# Paradoxical increases in anterior cingulate cortex activity during nitrous oxide-induced analgesia reveal a signature of pain affect

**DOI:** 10.1101/2023.04.03.534475

**Authors:** Jarret AP Weinrich, Cindy D Liu, Madison E Jewell, Christopher R Andolina, Mollie X Bernstein, Jorge Benitez, Sian Rodriguez-Rosado, Joao M Braz, Mervyn Maze, Mikhail I Nemenov, Allan I Basbaum

## Abstract

The general consensus is that increases in neuronal activity in the anterior cingulate cortex (ACC) contribute to pain’s negative affect. Here, using *in vivo* imaging of neuronal calcium dynamics in mice, we report that nitrous oxide, a general anesthetic that reduces pain affect, paradoxically, increases ACC spontaneous activity. As expected, a noxious stimulus also increased ACC activity. However, as nitrous oxide increases baseline activity, the relative change in activity from pre-stimulus baseline was significantly less than the change in the absence of the general anesthetic. We suggest that this relative change in activity represents a neural signature of the affective pain experience. Furthermore, this signature of pain persists under general anesthesia induced by isoflurane, at concentrations in which the mouse is unresponsive. We suggest that this signature underlies the phenomenon of connected consciousness, in which use of the isolated forelimb technique revealed that pain percepts can persist in anesthetized patients.

---

General anesthetics are potent regulators of pain processing and can produce effects ranging from diminished or absent pain perception (i.e., analgesia^1–4^) to the total abolition of reflexive responses to ongoing surgical stimuli^2, 5^. Importantly, pain is a conscious, multidimensional percept that includes both sensory-discriminative (modality, location, intensity) and affective-motivational (unpleasantness) features^6^. Studies in awake, responsive patients inhaling nitrous oxide, a general anesthetic gas with analgesic properties^1, 7, 8^, report a preferential reduction of the affective-motivational aspects of pain^9,10^. Currently, however, there is limited information as to the neural mechanisms that underlie the ability of nitrous oxide to modulate affective-motivational aspects of pain.

The anterior cingulate cortex (ACC) is critical to processing the affective-motivational features of the pain percept^11^. In humans, primates, and rodents, ACC neurons respond to noxious, but not innocuous, thermal and mechanical stimuli^12–15^. Human neuroimaging studies have shown that increased ratings of the unpleasantness of pain correlate with increased ACC activity, and that analgesia correlates with decreased ACC activity^11, 16^. Interestingly, after targeted ACC manipulations, including ablations^17–19^ or deep brain stimulation^20, 21^, patients still sense noxious stimuli, but report that the stimuli are less painful or less “bothersome”. Consistent with clinical findings, ablative^22, 23^ or pharmacological^24–26^ manipulations of the ACC in rodents produced selective reduction in affective-motivational responses to pain without influencing sensory thresholds^22–32^. Importantly, these studies led to the conclusion that *inhibition*^22–24^ of the ACC is critical to generating pain control and that *excitation*^25, 33–35^ produces increased pain aversion.

Unexpectedly, and indeed paradoxically given the general consensus^6, 36^, cerebral blood flow and metabolism-based measurements of neural activity report that nitrous oxide increases activity in frontal cortical regions^37^, and in particular in the ACC^38^. However, because nitrous oxide has a strong vasodilatory effect that confounds interpretations of data from neuroimaging studies^39, 40^, these findings are controversial. If true, these studies would suggest that increases in ACC activity do not necessarily lead to increased pain aversion and may even contribute to the analgesic effects of nitrous oxide. To date, however, there have been no direct measurements (electrophysiology or *in vivo* calcium imaging) of neural activity in the ACC during inhalation of nitrous oxide.

In this present work, using *in vivo* imaging of the calcium dynamics of ACC neurons in mice, we report that nitrous oxide, in fact, produces profound increases in ACC activity. Furthermore, in studies of molecularly distinct subsets of cortical neurons, we discovered that nitrous oxide preferentially activates excitatory ACC neurons, with limited actions on inhibitory interneurons. In behavioral studies, we confirmed that nitrous oxide produces a potent analgesia that preferentially diminishes affective-motivational pain endpoints. In awake, freely moving mice, we demonstrated that by increasing spontaneous ACC activity, nitrous oxide reduces the relative magnitude of noxious stimulus-induced ACC activation. Importantly, this reduction in noxious stimulus-evoked ACC activation correlates with nitrous oxide-induced reductions in affective-motivational behaviors, but not reflexive behaviors. Lastly, using these changes in ACC as a neural biomarker for affective-motivational aspects of pain, we demonstrate the presence of neural signatures of pain even in an isoflurane-anesthetized, behaviorally unresponsive mouse.

## RESULTS

### Nitrous oxide induces paradoxical increases in ACC activity

We used head-mounted miniature microscopes^41, 42^ to monitor the calcium dynamics^43^ of individually identified ACC neurons during exposure to nitrous oxide or control gas (oxygen) (**Fig. 1A, B**, **Fig. S1**). Across the two separate exposures, we identified 1,364 neurons (nitrous oxide: 795, oxygen: 569). Consistently, we found that inhalation of nitrous oxide drove sustained and significant increases in spontaneous ACC activity (**Fig. 1C**, **Supplemental Video 1**).

**Fig. 1.**
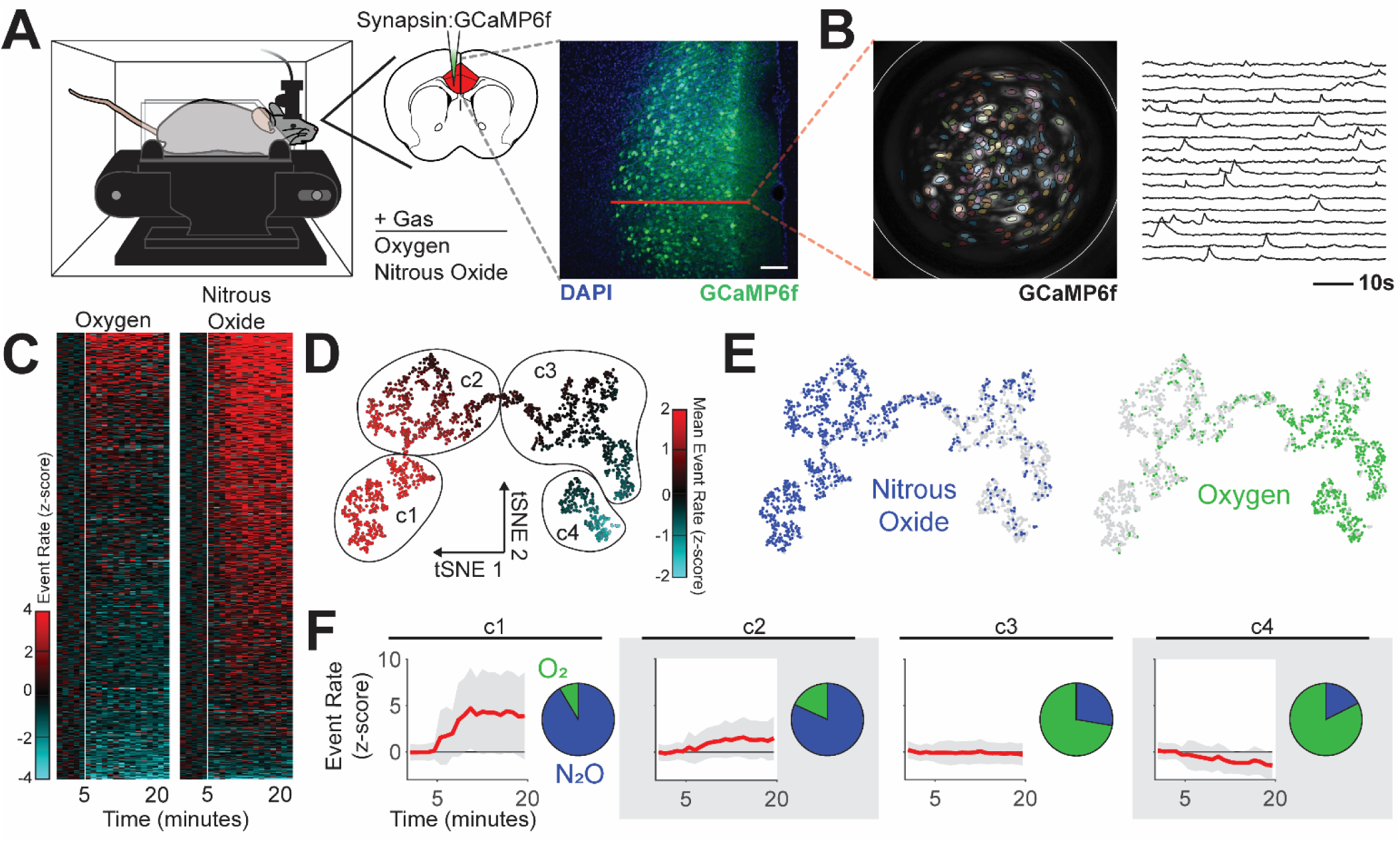
Nitrous oxide increases ACC activity. (**A**) Left: Spontaneous ACC activity monitored during inhalation of nitrous oxide using a genetically encoded calcium indicator (GCaMP6f) and head-mounted miniature microscopes. Right: ACC targeted GCaMP6f neuronal expression. Red line indicates target for calcium imaging. Scale bar (white) equals 50 µm. (**B**) Left: Maximum projection of recording over time displays neuronal distribution and is overlaid with PCA/ICA cell segmentation (colored areas) and GRIN lens boundaries (white circle). Right: Single neuron traces of extracted calcium fluorescence over time. (**C**) Heatmap of changes in event rate induced by nitrous oxide or control gas (oxygen); z-score normalized to baseline activity, prior to gas exposure (white line). (**D** and **E**) Representation of neural activity patterns using t-distributed stochastic neighbor embedding (tSNE), colored by neural activity (**D**) and gas exposure (**E**). (**F**) Identified clusters, including the mean (red) and standard deviation (gray) of neural activity (baseline normalized z-scored event rate) and cluster makeup by gas exposure.

Next, we used clustering to identify distinct activity patterns that occur during inhalation of nitrous oxide or control gas. Dimensionality reduction of neural activity patterns using t-distributed stochastic neighbor embedding (tSNE) reveals that neural activity patterns during inhalation of nitrous oxide are highly divergent and largely nonoverlapping with those observed during inhalation of control gas (**Fig. 1D** and **E**). Density-based spatial clustering after tSNE analysis identified 4 unique clusters of activity (**Fig. 1D** and **F**). Neurons identified during nitrous oxide recordings predominately populated clusters defined by large increases in activity (**Fig. 1F**: clusters 1, 2). Neurons from control gas recordings are largely confined to clusters with minimal activity changes (**Fig. 1F**: cluster 3) or slightly decreased activity (**Fig. 1F**: cluster 4). Thus, in sharp contrast to the established literature, we conclude that increased ACC activity can occur during inhalation of a known analgesic.

### Nitrous oxide preferentially activates excitatory ACC neurons in cortical layer 2/3

Cortical circuits are comprised of functionally distinct neurons with unique molecular identities that underlie cortical information processing^44–51^. Using a combinatorial viral/genetic strategy that enables the selective expression of genetically encoded calcium indicators within specific cell types, we monitored the activity of principal (excitatory) neurons (vGluT2 expressing; VG2), and molecularly distinct subpopulations of cortical inhibitory interneurons that express parvalbumin (PV), somatostatin (SST), or vasoactive intestinal peptide (VIP) (**Fig. 2A, B**). During separate exposures to nitrous oxide or control gas (oxygen) (**Fig. S1**), we identified 1,771 neurons (by subtype: VG2: 846, PV: 296, SST: 260, VIP: 369; by gas: nitrous oxide: 1049, oxygen: 722).

**Fig. 2.**
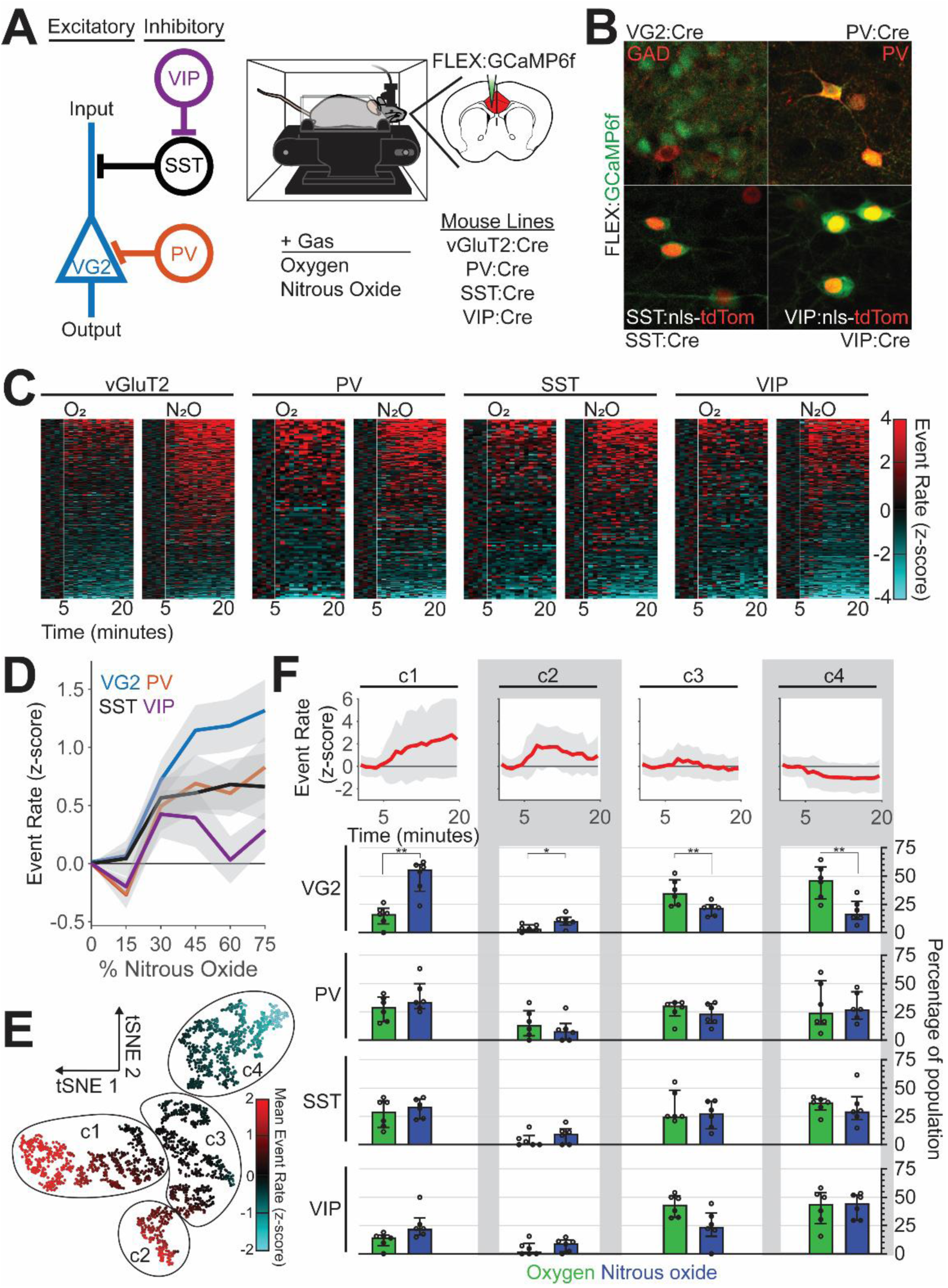
Nitrous oxide preferentially activates excitatory ACC neurons. (**A**) Left: Simplified circuit illustrating the local connectivity of molecularly distinct cortical neurons. Right: Spontaneous activity of molecularly distinct ACC neurons monitored during inhalation of nitrous oxide, after their restricted labeling with GCaMP6f using a combinatorial viral/genetic approach. (**B**) Selective labeling of molecularly distinct populations. (**C**) Heatmap of changes in event rate induced by nitrous oxide or control gas (oxygen); z-score normalized to baseline activity prior to gas exposure (white line). (**D**) Changes in neural activity (z-scored event rate) across different neural subtypes as a function of nitrous oxide concentration (colored line: mean, gray area: SEM). (**E**) Representation of neural activity patterns using t-distributed stochastic neighbor embedding (tSNE), colored by neural activity. (**F**) Mean normalized event rate of identified clusters (top; line: mean, gray area: standard deviation) and the preferential recruitment of distinct molecular subtypes to individual clusters by gas exposure, displayed as median and interquartile range (two-way repeated measures ANOVA, FDR corrected).

**Figures 2C** and **S2** show that compared to oxygen, nitrous oxide preferentially activated excitatory neurons. Although we do observe PV-, SST-, and VIP-expressing interneurons with increased activity (**Fig. 2D**), the proportion of neurons with increased activity did not differ between oxygen and nitrous oxide (**Fig. 2C** and **Fig. S2**). An assessment of the recruitment of subtypes of neurons to clusters with distinct activity profiles confirmed that excitatory, but not inhibitory (PV, SST, VIP), neurons are preferentially recruited to clusters with increased activity (**Fig. 2E** and **F**: clusters 1 and 2).

Next, to provide a surrogate of overall neuronal activity in the ACC, we immunolabeled for Fos protein (**Fig. 3A**), a molecular marker of recently activated neurons^52^. Paralleling our calcium imaging observations (**Fig. 1**), we found that nitrous oxide increased Fos expression compared to air (**Fig. 3B** and **C**). **Figure 3D** further shows that the increased Fos expression was particularly pronounced in cortical layers 2/3 compared to other layers. Also consistent with the recordings, we found a very small, albeit significant increase, from 3 to 6%, in Fos double-labeling of inhibitory interneurons (GAD67-GFP+). This result indicates that the increased Fos expression observed in layer 2/3 is largely due to activation of excitatory neurons. We conclude that nitrous oxide preferentially activates excitatory neurons in the ACC, effects that we hypothesize profoundly alter the experience of pain.

**Fig. 3.**
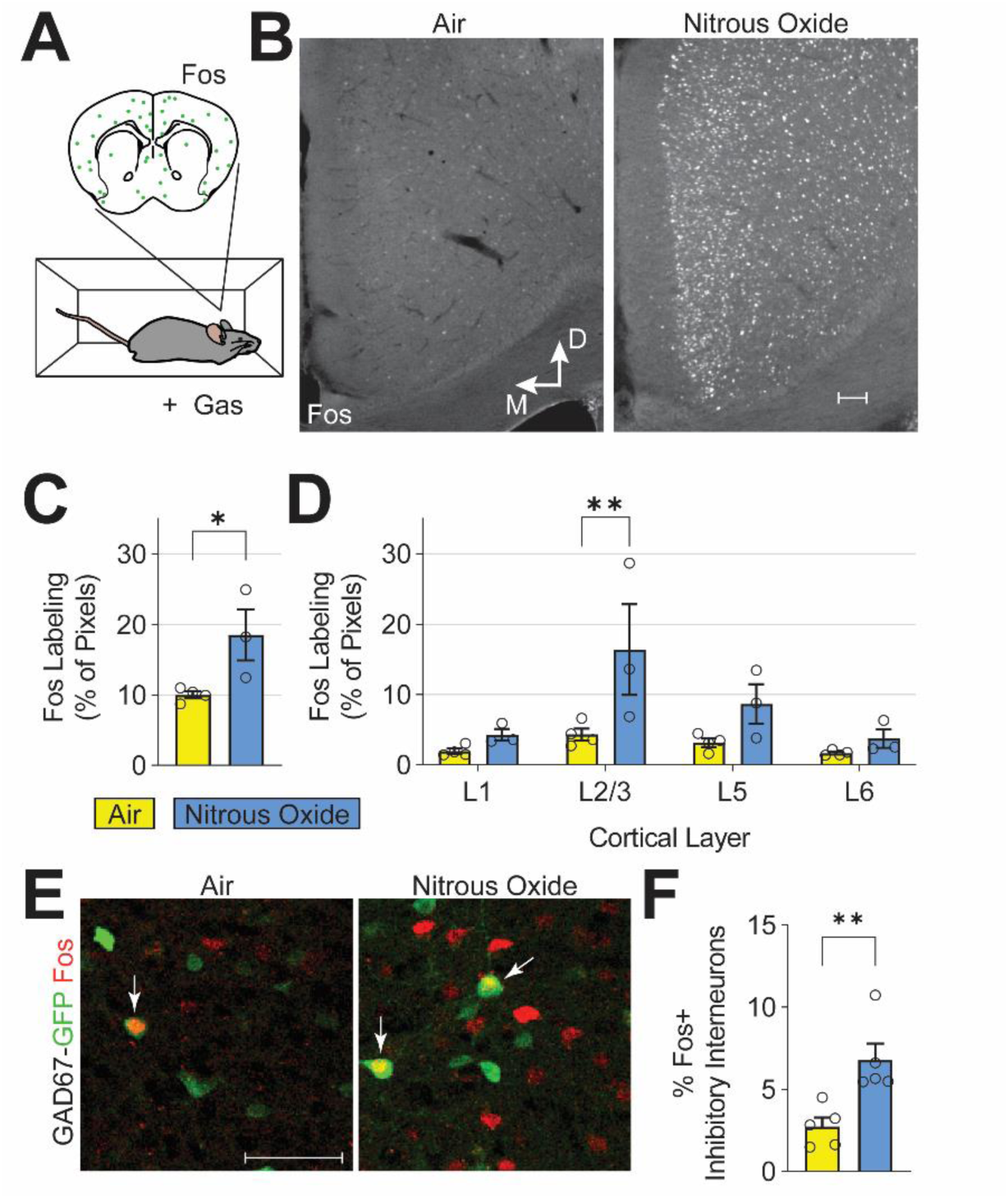
Nitrous oxide predominantly activates excitatory ACC neurons in cortical layer 2/3. (**A,B**) Fos immunofluorescence, a correlate of neuronal activity, after exposure to air or nitrous oxide (60%). (**C**) Quantification of ACC Fos labeling (Student’s t-test, p < 0.039). (**D**) Cortical layer-specific changes in Fos labeling (two-way ANOVA, FDR corrected). (**E**) Immunofluorescence labeling of Fos-expressing neurons (red) and inhibitory neurons with GFP in GAD67-GFP mice (green). White arrows indicate double-labeled cells. (**F**) Quantification of Fos-expressing ACC inhibitory interneurons (student’s t-test, p < 0.008). White scale bars in (**B**) and (**E**) equal 50 microns.

### Nitrous oxide reduces affective-motivational pain-related behaviors

Mice produce complex nocifensive behaviors to noxious stimuli. As noted above, alterations in ACC activity are postulated to influence affective-motivational, but not reflexive, responses to noxious stimuli. Here, we asked whether nitrous oxide-induced changes in ACC activity translate to preferential reduction of affective-motivational, rather than reflexive, indices of pain. In these studies, we compared the effects of nitrous oxide inhalation on the production of reflexive versus affective-motivational behaviors during presentation of a noxious stimulus (**Fig. 4A and B**).

**Fig. 4.**
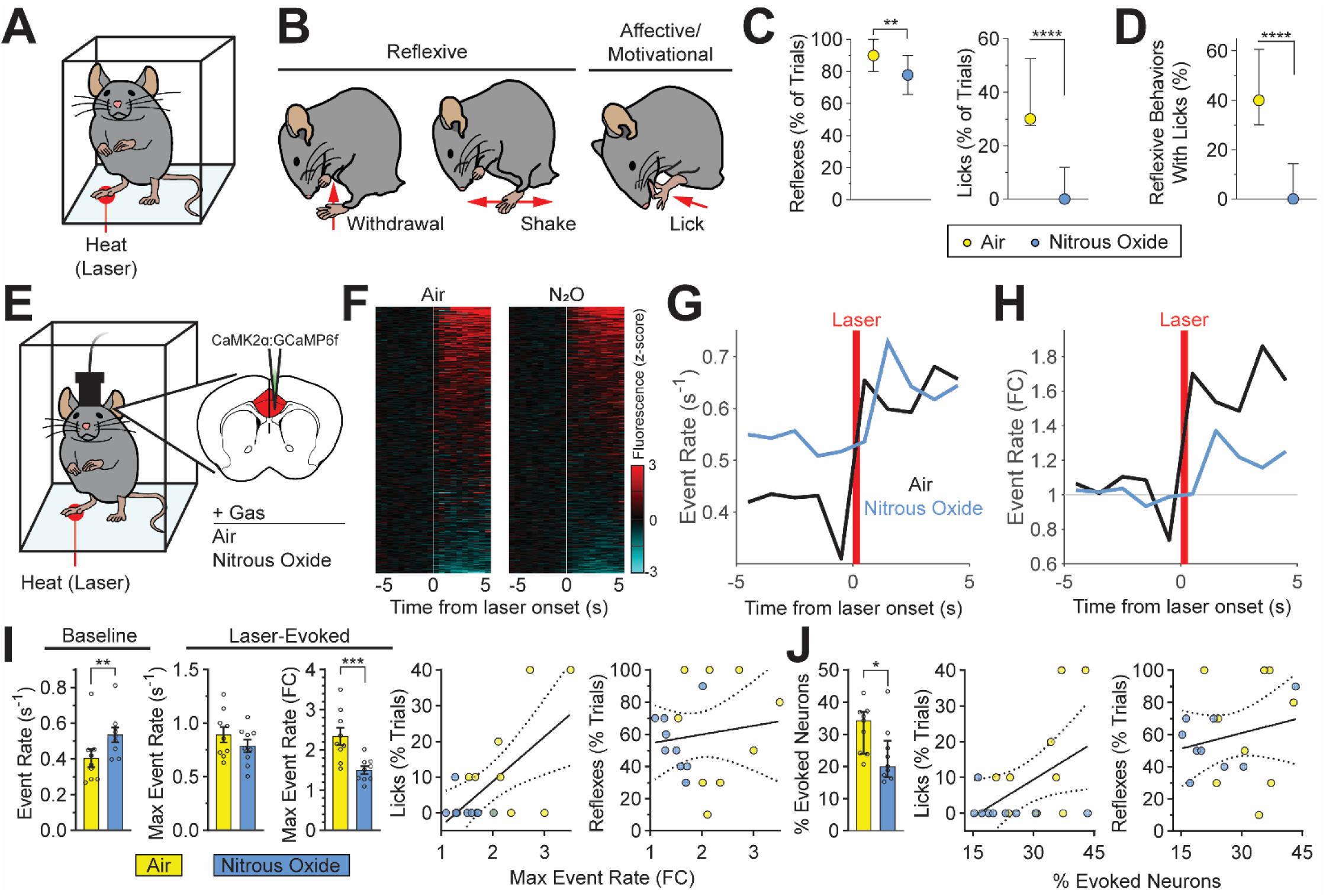
Nitrous oxide-induced reduction of affective-motivational pain-related behaviors correlates with changes in noxious stimulus-evoked ACC activity. (A) Behavioral responses to noxious heat (high-power infrared laser) monitored during inhalation of control gas (air) or nitrous oxide (60%). (B) Heat-evoked reflexive and affective-motivational behaviors. (C) Reflexive and affective-motivational behavioral responses to noxious stimuli during inhalation of nitrous oxide, quantified as percent of trials (paired t-test, reflexes: p < 0.0082; licks: p < 0.0001). (D) Percentage of licks that occur with reflexive behaviors (paired t-test, p < 0.0001). (E) Noxious stimulus-evoked ACC activity monitored in awake, freely behaving mice during inhalation of nitrous oxide. GCaMP6f virally-expressed in excitatory neurons (CaMK2α). (F) Heatmap of changes per neuron (rows) in calcium dynamics (z-scored and averaged across trials) provoked by laser stimulus (white line) during nitrous oxide or air. (G and H) Noxious stimulus-evoked neural activity during inhalation of nitrous oxide or air displayed as absolute event rate (G, events/second) and baseline normalized event rate (H) (n = 9 mice). (I) Left: Quantification of baseline and laser-evoked neural activity illustrated in G and H (paired t-test). Right: Simple linear regression of normalized maximum event rate versus licks (R^2^ = 0.382, p < 0.006) or reflexes (R^2^ = 0.017, p < 0.610). (J) Left: Neurons with significantly altered calcium dynamics following laser stimulation as a percentage of the total number of neurons identified per mouse, quantified from (F) (paired t-test). Right: Simple linear regression of the percentage of neurons with altered activity versus licks (R^2^ = 0.235, p < 0.042) or reflexes (R^2^ = 0.046, p < 0.390). Data for plots in C, D and J displayed as median and interquartile range; bar graphs in I displayed as mean ± SEM; regressions in I and J displayed as best fit line and 95% confidence interval. N = 30 mice for panels C and D. N = 9 mice for panels F through J.

While mice inhaled nitrous oxide (60%) or control gas (air), we generated brief, noxious heat stimuli using infrared laser pulses targeted to the hindpaw. The laser stimulus produced robust nocifensive responses in mice, including withdrawals, shakes, and licks^53^ (**Fig. 4B**, **Supplemental Video 2**). Importantly, in rodents, licking of the hindpaw following a noxious stimulus is an affective-motivational response indicative of the experience of pain^27, 54–56^, not merely a reflexive response, as is the case for withdrawals and shakes^55, 57, 58^. **Figure 4C** shows that nitrous oxide produces a potent suppression of affective-motivational measures of pain. Indeed, nitrous oxide almost completely abolished laser-evoked licks. In contrast, reflexive behaviors are minimally affected by nitrous oxide (**Fig. 4C**).

In mice, a single laser stimulus often evokes multiple concurrent behaviors, with the production of licks usually coupled to reflexive behaviors (i.e., withdrawals and shakes). Nitrous oxide dramatically reduced this coupling, with laser-evoked reflexes producing proportionally fewer concurrent licks (**Fig. 4D**). Taken together, we conclude that the reduction of affective-motivational behaviors produced by nitrous oxide is independent of any effects on reflexive behaviors.

### Nitrous oxide-induced analgesia correlates with noxious stimulus-evoked ACC activity

Next, we investigated how paradoxical, nitrous oxide-induced increases in spontaneous ACC activity translate to an analgesic effect in response to a noxious stimulus. We hypothesized that nitrous oxide-induced increases in ACC activity create a ceiling effect, thereby attenuating the relative magnitude of noxious stimulus-evoked changes, which results in a diminished experience of pain. To test this hypothesis, we imaged neural activity in ACC neurons during inhalation of nitrous oxide (60%) or control gas (air) with concurrent delivery of the laser stimulus (**Fig. 4E**, **Supplemental Video 3**). Given our finding that nitrous oxide predominantly increases activity in excitatory neurons of the ACC, here we used a viral approach to monitor activity exclusively in excitatory neurons (**Fig. 4E**). Across the nitrous oxide and air exposures, we identified 1402 neurons (nitrous oxide: 825, air: 577).

During presentation of the laser stimulus, we found that noxious stimulus responsive ACC neurons accounted for 31.8±2.5% of the total population in the air condition and 23.4±3.0% under nitrous oxide (**Fig. 4F, J**). In agreement with the findings described above, nitrous oxide significantly increased the pre-stimulus baseline event rate compared to air (**Fig. 4G** and **I**). Interestingly, and as would be expected for a ceiling effect, we observed no difference in the absolute magnitude of laser-evoked activity between nitrous oxide and air conditions (**Fig. 4G** and **I**). However, when measures of noxious stimulus-evoked activity were normalized to pre-stimulus baseline activity there was a clear reduction in the nitrous oxide exposure as compared to air (**Fig. 4H** and **I**). In other words, nitrous oxide reduces the relative magnitude of noxious stimulus-evoked activity compared to air.

Importantly, the degree of laser-evoked licking behavior, but not reflexes, correlates not only with the relative magnitude of the laser-evoked maximum event rate, but also with the percentage of ACC neurons activated by the laser (**Fig. 4I**, **J**). These relationships suggest that noxious stimulus-evoked neural activity of the ACC can indeed be used as a proxy for the affective-motivational aspects of the pain experience in mice.

### Neural signatures of pain are present during isoflurane-induced general anesthesia

To date, aside from directly asking patients to rate levels of ongoing pain, there is no neural biomarker that can be used as a proxy of the affective experience of pain. This lack of an adequate pain biomarker is particularly problematic during general anesthesia, where patients (and animals) are immobilized and thus behaviorally unresponsive and incapable of reporting their pain experience^59^. Rather disturbingly, clinical studies using the isolated forearm technique reveal that patients often report the experience pain under general anesthesia^60, 61^. Fortunately, perhaps, the amnestic effects of general anesthetics largely render them unable to recall such events postoperatively^62^. As nitrous oxide does not produce loss of behavioral responsiveness in mice under normal conditions (concentrations greater than 100% would be required), we could not determine whether there is persistent brain activity in a nitrous oxide anesthetized mouse. Therefore, we initiated studies using isoflurane, a widely used volatile anesthetic that readily produces general anesthesia. Our specific question is whether there can be persistent ACC activity consistent with the experience of pain in an otherwise anesthetized, behaviorally unresponsive mouse.

We first assessed the influence of isoflurane on the spontaneous activity of ACC neurons. In contrast to changes recorded during nitrous oxide inhalation, we found that isoflurane decreased spontaneous ACC activity in a dose-dependent manner, and completely abolished ACC activity at the highest concentrations (**Supplemental Video 4**).

We then assessed noxious stimulus-evoked ACC activity in isoflurane-anesthetized mice. As expected, mice tested at 2% isoflurane, a concentration where nociceptive reflexes withdrawals are abolished, we observed a complete absence of spontaneous and evoked activity.

Next, we tested mice during inhalation of 1% isoflurane in air, a concentration where mice are immobilized and lack righting reflexes (a rodent measure of awareness), yet still retain neural activity (**Fig. 5A** and **B**). Although laser stimulation increased the activity of ACC neurons in both air and 1% isoflurane conditions (**Fig. 5C**), we recorded significantly fewer laser responsive neurons under isoflurane (**Fig. 5G**). However, we found no significant differences between baseline or laser-evoked ACC activity (i.e., event rate-based) measurements (**Fig. 5E, F, G**).

**Fig. 5.**
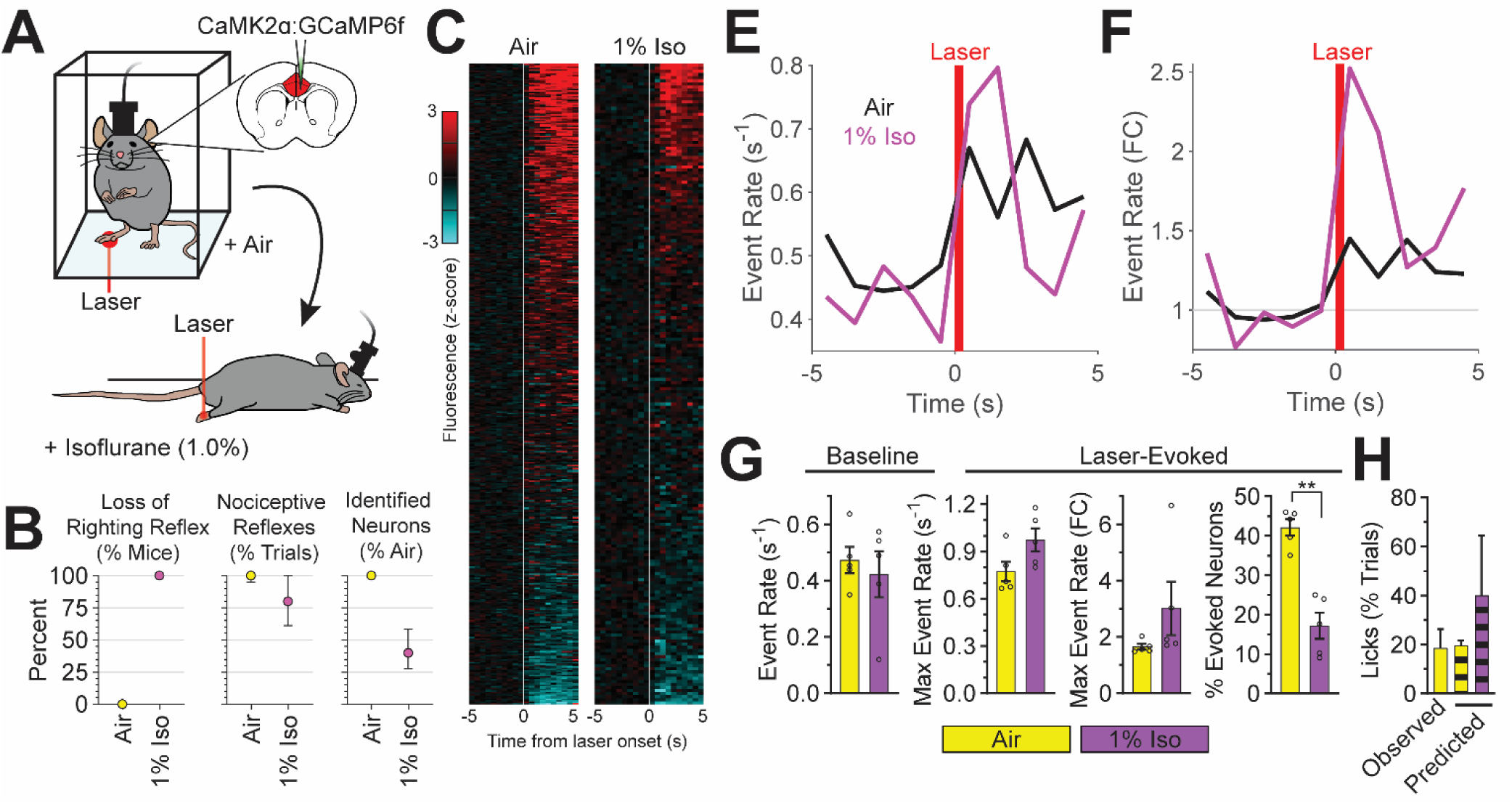
Neural signatures of pain during general anesthesia. (**A**) Noxious (laser) stimulus-evoked ACC activity monitored in awake, freely behaving mice inhaling air and in anesthetized mice inhaling isoflurane (1%). GCaMP6f virally-expressed in excitatory neurons (CaMK2α). (**B**) Behavioral and imaging endpoints: left: loss of righting reflex/absence of volitional movements; middle: presence of nociceptive reflexes; right: percent of spontaneously active neurons (isoflurane compared to air). Data displayed as median and interquartile range. (**C**) Heatmap of calcium dynamics (z-scored and averaged across trials) per neuron in response to laser stimulus (white line) during isoflurane or air. (**E** and **F**) Laser-evoked neural activity during inhalation of isoflurane or air displayed as absolute event rate (**E**, events/second) and baseline normalized event rate (**F**). (**G**) Baseline and laser-evoked neural activity quantified from **E** (baseline and max event rate), **F** (fold change in max event rate), and **C** (percent of neurons with evoked activity) (paired t-test). (**H**) Noxious stimulus-evoked affective-motivational behaviors (licks) observed in awake mice (solid bar) and those predicted by noxious stimulus-evoked ACC activity (striped bars). N = 5 mice per group for all panels.

Lastly, using the relationship between noxious stimulus-evoked ACC activity and the generation of affective-motivational behaviors (licks) recorded in nitrous oxide and air exposed mice (**Supplemental Fig. 3**), we assessed the presence of neural activity patterns indicative of an affective-motivational pain experience in isoflurane-anesthetized mice. Surprisingly, noxious stimulus-evoked ACC activity during inhalation of 1% isoflurane, when mice could not perform licks due to isoflurane-induced immobilization, not only persists, but is consistent with the experience of pain (**Fig. 5H**). We conclude that a brain signature of affective-motivational aspects of the pain experience can be preserved under general anesthesia.

## DISCUSSION

In this study, we explored the influence of nitrous oxide, an inhalational anesthetic with analgesic properties, on neural activity of the ACC, a cortical region that is a major contributor to the affective-motivational aspects of pain^6^. In contrast to the prevailing view, namely that inhibition of the ACC reduces pain affect^22–24^, we discovered that nitrous oxide profoundly increases spontaneous ACC neuronal activity. Clearly, our results present a paradox: how is it that nitrous oxide increases spontaneous ACC activity and produces analgesia, but does not, as the literature would predict, increase affective-motivational indices of pain. Unexpectedly, we discovered that the absolute magnitude ACC activity provoked by a noxious stimulus did not differ between nitrous oxide and air, even though nitrous oxide reduced measures of the affective pain experience. Rather, we demonstrate that it is the relative magnitude of noxious stimulus-evoked ACC activity, as compared to activity immediately prior to the stimulus, that best correlates with the production of affective-motivational pain behaviors. In essence, what underlies the affective-motivational aspects of pain by the ACC is a circuit mechanism of gain control^63^, namely one that adjusts the signal-to-noise ratio of stimulus-evoked ACC neuronal activity^64^. In this model, spontaneous activity (i.e., noise) can be tuned, for example, by nitrous oxide, to modulate the relative change of activity provoked by a noxious stimulus (i.e., signal). Based on our findings, we conclude that increased activity *per se* is not necessarily indicative of a pain experience. Rather, it is the change in between resting (spontaneous) ACC activity and that evoked by a noxious stimulus (e.g., laser or surgical intervention) that determines whether there is pain affect. Of course, this conclusion is consistent with the fact that despite ongoing activity in the naïve mouse (and human), there is no pain affect until the introduction of a noxious stimulus.

Unclear is the mechanism underlying the selective increase in the activity of excitatory ACC neurons by nitrous oxide, purportedly a non-competitive inhibitor of the NMDA receptor^65^. Although one would expect that blocking NMDA receptors would decrease neuronal activity, previous recordings in the prefrontal cortex reported that selective NMDA receptor antagonists, in fact, increase excitatory neuronal activity, not directly, but by decreasing the activity of inhibitory interneurons^66^. As we did not observe decreases in inhibitory interneuron activity, we suggest that nitrous oxide’s effects on the ACC involve alternative mechanisms^65^. For example, nitrous oxide could influence ACC neuronal activity via direct actions on upstream brain regions^67^, such as medial-dorsal thalamus or basolateral amygdala^68^.

Also unclear are the direct downstream consequences of nitrous oxide-induced increases in ACC activity, and how this translates to behavioral analgesia in tests of pain affect. The ACC is a major hub that is highly connected to other elements of the so-called “pain matrix”^69^. Thus, nitrous oxide-induced activation of ACC projection neurons would produce wide-ranging effects on other components of the matrix^70^, thereby altering the experience of pain. Other studies reported that nitrous oxide analgesia is naloxone reversible^65^. In ongoing studies we are examining whether the downstream circuits engaged by the ACC contribute to the naloxone-reversible aspects of nitrous oxide-induced analgesia, potentially by direct actions on endorphin-mediated inhibitory controls^71, 72^.

Particularly surprising was the persistence of noxious stimulus-evoked activity in the ACC of isoflurane-anesthetized mice, at concentrations that blocked behavioral indices of pain affect, namely, licking in response to a noxious stimulus. As this activity was comparable to that recorded in awake mice, we suggest that it represents a neural biomarker, in effect a surrogate pain index that is specific for the affective component of the pain experience. In other words, the brain can “experience” pain even under general anesthesia, an interpretation consistent with provocative clinical reports of the high prevalence of an intraoperative experience of pain in patients^60–62, 73^ under general anesthesia. These patients can communicate their pain experience in real time through the isolated forearm technique^59–61^, a phenomenon known as connected consciousness^59^.

Importantly, it is possible to establish a level of general anesthesia in which connected consciousness does not occur^74^. In fact, when we increased the depth of anesthesia using 2% isoflurane and then tested the mice, not only was there no behavioral response to a noxious stimulus, but we observed that both spontaneous and evoked ACC activity were abolished. This absence of activity at the deepest levels of anesthesia^75^ creates, in essence, a functional “ablation” of the ACC, which blocks the experience of pain much in the same way that a physical lesion of the ACC provides pain relief in patients^17^. However, although similarly deep levels of general anesthesia (measured by EEG) can ensure the absence of connected consciousness^74^, the associated increased risk of adverse postoperative outcomes (death, stroke, postoperative delirium)^76^ likely outweigh any potential benefits. For this reason, our findings are particularly relevant to ongoing efforts to develop neural activity-based biomarkers that can reliably document adequate analgesia during surgery under general anesthesia^77^, which, in turn, will support the development of novel general anesthetics that can safely block the experience of pain.

## METHODS

### Animal husbandry

All mouse husbandry and surgical procedures adhered to the regulatory standards of the Institutional Animal Care and Use Committee of the University of California San Francisco (UCSF; protocol AN199730). The following mouse strains were used: vGluT2-IRES-Cre^78^ (Jax # 028863), PV-IRES-Cre^79^ (Jax #017320), SST-IRES-Cre^80^ (Jax # 028864), VIP-IRES-Cre^80^ (Jax # 031628), Ai75D (ROSA26-nls-tdTomato; Jax # 025106), and GAD67-GFP^81^. The health and wellbeing of the mice were monitored daily.

### Calcium imaging of spontaneous ACC activity

#### GECI expression strategy

We used two strategies to express genetically encoded calcium indicators (GECI; GCaMP6f^43^) in neurons: (1) viral pan-neuronal expression, and (2) viral/genetic expression within molecular distinct subsets of neurons. For experiments with pan-neuronal expression, we delivered GCaMP6f under control of the synapsin promoter (AAV1/9-SYN-GCaMP6f; Addgene, #100837)^51^. The restricted delivery of GCaMP6f to molecularly distinct populations of neurons was achieved using the vGluT2-Cre^78^, PV-Cre^79^, SST-Cre^80^, or VIP-Cre^80^ mouse lines in combination with the Cre-inducible viral expression of GCaMP6f in the ACC (AAV1/9-SYN-FLEX-GCaMP6f; Addgene, #100833).

#### Surgical preparation for ACC calcium imaging

Briefly, mice were anesthetized with isoflurane (2% in oxygen) and placed on a stereotaxic frame (Kopf). After craniotomy above the left ACC (Bregma, x: -0.33mm, y:1.27mm), we injected virus (depth: -1.75mm), and chronically implanted a gradient index (GRIN) lens (0.5×4mm ProView, Inscopix; depth: -1.7mm). The GRIN lens and titanium headbar (custom made, eMachineShop.com) were affixed to the skull with dental cement (Metabond). Mice were provided with postoperative analgesia (carprofen and slow-release buprenorphine). One week after implantation surgery, under isoflurane anesthesia, a baseplate was affixed above the GRIN lens with dental cement. To provide time for sufficient GCaMP6f expression, mice recovered for 3 to 4 weeks before experiments began.

#### Behavioral apparatus and anesthesia delivery

Mice were headfixed to a passive treadmill^82^, which was modified to provide heating that kept the mice isothermic during exposure to anesthesia, and then placed in a modified anesthetic induction chamber (VetEquip, 7L). Before the start of the experiment, the atmosphere of the chamber was replaced with oxygen. During experimental sessions, the mice were exposed to continually increasing concentrations of isoflurane or nitrous oxide, or for control conditions, continued exposure to oxygen. For all experiments, the concentration of oxygen never fell below 21%. Isoflurane was delivered via an Isoflurane Vaporizer (DRE Veterinary). Gas concentrations were monitored by a Datex Ohmeda S/5 anesthesia patient monitor and recorded by VSCapture software.

#### Calcium imaging and behavior monitoring

Changes in GCaMP6f fluorescence were captured with Inscopix miniscopes (nVista 3.0 or nVoke 2.0) at 20 frames per second (fps). Imaging parameters (excitation LED power, digital gain, and focus depth) were individually set for each mouse. Calcium imaging data were recorded via Inscopix Data Acquisition Software (IDAS), and recordings were triggered via TTL input. The behavior of the mouse was monitored with a Logitech webcam and recorded via ffmpeg software (https://ffmpeg.org/). Recording sessions were coordinated by Arduino/MATLAB, which triggered the start and end of data acquisition via miniscopes, ffmpeg, and VSCapture.

#### Tissue processing

After completion of the *in vivo* imaging experiments, the mice were anesthetized with Avertin (2.5% in saline) and transcardially perfused with phosphate-buffered saline (PBS) and then 4% formaldehyde (37% formaldehyde; Acros Organics, 11969-0100) diluted in PBS. Whole heads were postfixed in 4% formaldehyde at 4°C overnight, then brains were extracted from the skull and postfixed overnight at 4°C. Following postfixation, the brains were cryoprotected at 4°C overnight in 30% sucrose in PBS, and embedded in specimen matrix (Optimal Cutting Temperature (OCT) compound, Tissue-Tek) and stored at -80°C. Confocal microscopy confirmed GCaMP6f expression and proper GRIN lens targeting.

### Assessing induction of immediate early genes by nitrous oxide

#### Fos induction

Adult GAD67-GFP mice (6-10 weeks old) were habituated to an anesthetic induction chamber (7L, VetEquip) for 30-minute sessions on 3 separate days. The following day, after an additional 30 minute habituation, the mice were exposed to 2L/minute of 60% nitrous oxide or medical air. After 2 hours, the mice were anesthetized with Avertin and transcardially perfused as described above. The brain was then removed, post-fixed in 4% formaldehyde for 4 hours at 4°C and then cryoprotected in 30% sucrose, embedded in OCT, frozen on dry ice, and stored at -80°C.

#### Fos immunohistochemistry and confocal imaging

Frozen brains were coronally sectioned (30 microns) with a Hacker cryostat (Bright OTF series). ACC sections were slide mounted, washed with PBS (3 times for 5 minutes), and blocked with 10% normal goat serum (NGS) in PBS for 1 hour. Slides were incubated overnight at room temperature in rabbit anti-Fos primary antibody (1:1000, Cell Signaling Technology) diluted in PBS with 0.3% Triton-X and 1% NGS (PBST). Slides were then washed with PBS (3 times for 5 minutes), incubated in AlexaFluor 594-conjugated goat anti-rabbit secondary antibody (1:1000, Invitrogen) diluted in PBST at room temperature for one hour, and washed again with PBS (3 times for 5 minutes). Sections were coated with mounting media (DAPI Fluoromount-G, Southern Biotech, #0100-20), and then coverslipped with #1.5 glass (Epredia, #152460). Fos immunofluorescence was captured via epifluorescent microscopy using a Zeiss Axio Zoom.V16. Colocalization of GAD67-GFP and Fos was captured by confocal microscopy using a Zeiss LSM980 Airyscan II microscope.

#### Fos image analysis and quantification

Fos expression was quantified using a thresholded z-scoring approach with custom-written MATLAB code. Briefly, we used the intensity values of pixels in cortical layer 1, which has minimal Fos expression, to set the mean and standard deviation for image z-scoring. The intensity of all pixels within an image were z-scored as: (Pixel intensity – mean background intensity)/(standard deviation of background intensity). Pixels within the ACC that have z-score values above 1.96 were considered Fos+. Double labeling of GAD67-GFP and Fos immunolabeling was quantified within ImageJ by a blind scorer.

### Assessing noxious stimulus-evoked responses to infrared laser pulses

#### Behavioral apparatus and volatile anesthetic delivery

Mice were placed inside a modified anesthetic induction chamber (VetEquip, 2L) with a high-transmittance glass floor that allowed for the presentation of noxious heat stimuli during the concurrent inhalation of nitrous oxide. During experimental sessions, mice were exposed to nitrous oxide (60%) or control gas (medical air). We used subhypnotic concentrations of nitrous oxide that allowed awake, weightbearing mice to freely respond to noxious stimuli^83^, but below concentrations that induce unconsciousness (i.e., MACawake)^84, 85^. The concentration of oxygen was held equivalent to atmospheric concentrations (21%) during nitrous oxide inhalation. Concentrations of nitrous oxide, oxygen, and carbon dioxide were monitored by a Datex Ohmeda S/5 anesthesia patient monitor and recorded using VSCapture software^86^. Individual mice were tested with different gasses during separate experimental sessions using a crossover study design with a minimum washout period of 7 days.

#### Generation of acute noxious thermal stimuli

Acute noxious thermal stimuli were generated using a fiber-attached infrared diode laser (LASMED (Lass-7M) 7W 975nm laser) that produced brief pulses that rapidly heat skin without causing injury^53, 87^. Mice received 10 trials of laser stimuli, with one laser pulse per trial. The laser power and pulse duration were set to 1750mA and 300 milliseconds, respectively. During presentation of the laser stimulus, a focused beam (2.0mm, 1/e^2^ diameter) was shone on the central portion of the plantar surface of the hindpaw^53^. The laser stimulus was manually triggered via a footswitch and laser firing time was recorded via Arduino/MATLAB. Behavioral responses were recorded with a digital camera (Imaging Source, DMK 37BUX252) at 200 frames per second (fps) using StreamPix (Norpix) software.

### Calcium imaging of noxious stimulus-evoked ACC activity

#### ACC calcium imaging preparation

Mice were prepared for calcium imaging as described above, with the following differences: (1) GCaMP6f was virally expressed in excitatory neurons using the CaMKIIa promotor (AAV1/9-CaMKIIa-GCaMP6f, Inscopix), (2) viral injection and implantation of an integrated GRIN lens/baseplate (0.5×4mm, Inscopix) occurred within a single surgery.

#### Calcium imaging and behavior monitoring

Changes in GCaMP6f fluorescence were monitored and noxious heat stimuli were generated as described above. For nitrous oxide recordings, anesthesia and behavioral monitoring were performed as for the laser experiments. For recordings under isoflurane anesthesia, mice were first tested during inhalation of air as described above. Then, mice were briefly anesthetized with 1.0% isoflurane within the recording chamber, and then moved from the chamber to a heating pad where isoflurane was administered via nosecone. Mice were equilibrated to isoflurane anesthesia for 30 minutes to ensure steady-state concentrations within the brain^88^ before testing resumed. This equilibration process was repeated for recordings conducted under 2% isoflurane. Miniscope recording, laser pulses and behavioral camera recording signals were coordinated via TTL pulse via Arduino/MATLAB and synchronized by monitoring TTL signals via Inscopix DAQ (data acquisition) box.

### Data processing

#### Calcium imaging data processing

Calcium imaging data were processed with Inscopix Data Processing Software (IDPS), MATLAB-IDPS API, and custom MATLAB code. Briefly, raw videos are spatially cropped, downsampled (2X), and bandpass filtered^89^. Processed videos are then motion corrected, normalized (dF/F), and individual cells segmented using principal component analysis and independent component analysis (PCA/ICA)^90^. Cell segmentation is manually confirmed in the IDPS GIU for each identified cell. Changes in calcium fluorescence per cell are extracted, and individual calcium transients within a trace are extracted as events, using either event detection algorithm in IDPS or custom MATLAB code.

#### Calcium imaging data analysis

To demonstrate nitrous oxide-induced increases in spontaneous activity, we Z-score normalized the activity of each cell to the baseline event rate. For clustering analysis, z-scored neuronal activity was transformed by dimensional reduction using tSNE algorithm. Clusters of neurons with unique activity patterns were identified from the tSNE mapping using DBSCAN. For stimulus-evoked changes, activity was extracted from 10 separate presentations of the noxious laser stimulus. The timing of stimulus presentation was determined using laser-generated TTL pulses recorded via the Inscopix nVoke DAQ box. Pre-stimulus activity is the time from -5 to 0 seconds before the laser stimulus; post-stimulus activity is measured from 0 seconds to 5 seconds after the laser stimulus. Baseline event rate is the mean pre-stimulus event rate per neuron per mouse. The maximum event rate is the maximum post-stimulus event rate. Where noted, fold changes in measures of neural activity are calculated as post-stimulus activity divided by pre-stimulus activity. Fluorescence traces were used to calculate the percentage of neurons with stimulus-evoked activity. Traces were z-scored to pre-stimulus activity and considered to have evoked activity if the trace z-scored value was greater than 1.96 for the post-stimulus period. Stepwise linear regression with interaction effects was performed on data mean-centered to control gas condition (medical air only). A neural activity (biomarker)-based estimate of laser evoked licks that otherwise would occur in awake mice was predicted from ACC recordings in isoflurane anesthetized mice. The isoflurane data were mean-centered to the control gas (medical air), and a prediction was made using the linear regression model generated from nitrous oxide and air recordings.

#### Scoring of stimulus-evoked behaviors

We used custom MATLAB software to score behavioral videos (viewed at 1/5X speed). Behavioral responses to individual laser stimuli were categorized as: (a) no response, (b) reflexes, such as flinches (the stimulated hindpaw shifted position but did not leave the glass floor), withdrawals (the stimulated hindpaw is rapidly pulled off of the glass floor), or shakes (the stimulated hindpaw is moved in a repetitive oscillatory fashion), or (c) licks (the stimulated hindpaw is brought to the face and licked or bitten). Each video was scored independently by two individuals. All scorers were blind to experimental conditions.

### Statistical analyses

Data were processed and analyzed in Mathwork’s MATLAB (R2020a) software. Statistical tests were performed with Prism (GraphPad) software. The threshold for significance for all statistical tests was set at p < 0.05, and indicators of significance levels were as follows: ns (not significant; p > 0.05); *= p < 0.05; **=p < 0.01; *** =p < 0.001; and ****= p < 0.0001. Corrections for multiple comparisons were performed using the false discovery rate method of Benjamini, Krieger, and Yekutieli, and noted within figure legends as “FDR corrected”.

## ACKNOWLEDGEMENTS

This work was funded by NIH NSR35NS097306 (AIB), Open Philanthropy (AIB), NIGMS fellowship F32GM131479 (JAPW), and Canadian Institutes of Health Research (CIHR) Doctoral Foreign Study Award DFD-170771 (CDL). We are grateful to: Dr. Eric Lam (UCSF) for aiding in the design and fabrication of experimental setups; Dr. Philip Bickler (UCSF) for advice on experimental setup; Dr. Racheli Wercberger (UCSF) for helpful advice on calcium imaging and analyses; Dr. Chris Tsang (Inscopix) for initial training for surgical implant of GRIN lenses; Dr. Julian Motzkin (UCSF) for helpful discussions on data analysis and statistics.

## SUPPLEMENTAL VIDEOS

**Supplemental Video 1. Nitrous oxide-induced changes in pan-neuronal ACC activity.** *In vivo* calcium imaging of ACC activity during inhalation of increasing concentrations of nitrous oxide. GCaMP6f fluorescence is normalized (dF/F). Video is played at 10X speed.

**Supplemental Video 2. Behavioral responses to laser stimuli.** Representative videos of lack of response, or laser-evoked responses (withdrawal, shake, lick).

**Supplemental Video 3. Laser-evoked ACC activity with simultaneous behavior monitoring.** Left: Front view of mouse in anesthesia chamber during miniscope recording with concurrent laser stimulation. Middle: Bottom view of mouse during laser stimulus. Right: Laser-evoked activity in the ACC (dF/F normalized).

**Supplemental Video 4. Isoflurane-induced changes in pan-neuronal ACC activity.** *In vivo* calcium imaging of ACC activity during inhalation of increasing concentrations of isoflurane. GCaMP6f fluorescence is normalized (dF/F). Video is played at 10X speed.

## REFERENCES

1. Davy, H. Researches, Chemical and Philosophical; Chiefly Concerning Nitrous Oxide: Or Dephlogisticated Nitrous Air, and Its Respiration. 588 (1800).

2. Snow, J. D. On the inhalation of the vapour of ether in surgical operations: Part I. in On the inhalation of the vapour of ether in surgical operations (John Churchill, 1847).

3. Coffey, F. et al. STOP!: a randomised, double-blind, placebo-controlled study of the efficacy and safety of methoxyflurane for the treatment of acute pain. Emerg. Med. J. 31, 613–618 (2014).

4. Olofsen, E. et al. Ketamine psychedelic and antinociceptive effects are connected. Anesthesiology 136, 792–801 (2022).

5. Merkel, G. & Eger, E. I. A comparative study of halothane and halopropane anesthesia including method for determining equipotency. Anesthesiology 24, 346–357 (1963).

6. Basbaum, A. I., Bautista, D. M., Scherrer, G. & Julius, D. Cellular and molecular mechanisms of pain. Cell 139, 267–284 (2009).

7. Sanders, R. D., Weimann, J. & Maze, M. Biologic effects of nitrous oxide: a mechanistic and toxicologic review. Anesthesiology 109, 707–722 (2008).

8. Lew, V., McKay, E. & Maze, M. Past, present, and future of nitrous oxide. Br. Med. Bull. **125**, 103–119 (2018).

9. Pirec, V., Patterson, T. H., Thapar, P., Apfelbaum, J. L. & Zacny, J. P. Effects of subanesthetic concentrations of nitrous oxide on cold-pressor pain in humans. Pharmacol. Biochem. Behav. 51, 323–329 (1995).

10. Ramsay, D. S., Brown, A. C. & Woods, S. C. Acute tolerance to nitrous oxide in humans. Pain 51, 367–373 (1992).

11. Rainville, P., Duncan, G. H., Price, D. D., Carrier, B. & Bushnell, M. C. Pain affect encoded in human anterior cingulate but not somatosensory cortex. Science 277, 968– 971 (1997).

12. Hutchison, W. D., Davis, K. D., Lozano, A. M., Tasker, R. R. & Dostrovsky, J. O. Pain-related neurons in the human cingulate cortex. Nat. Neurosci. 2, 403–405 (1999).

13. Kuo, C.-C. & Yen, C.-T. Comparison of anterior cingulate and primary somatosensory neuronal responses to noxious laser-heat stimuli in conscious, behaving rats. J. Neurophysiol. 94, 1825–1836 (2005).

14. Iwata, K. et al. Anterior cingulate cortical neuronal activity during perception of noxious thermal stimuli in monkeys. J. Neurophysiol. 94, 1980–1991 (2005).

15. Kuo, C.-C., Chiou, R.-J., Liang, K.-C. & Yen, C.-T. Differential involvement of the anterior cingulate and primary sensorimotor cortices in sensory and affective functions of pain. J. Neurophysiol. 101, 1201–1210 (2009).

16. Kröger, I. L. & May, A. Central effects of acetylsalicylic acid on trigeminal-nociceptive stimuli. J. Headache Pain 15, 59 (2014).

17. Foltz, E. L. & White, L. E. Pain “relief” by frontal cingulumotomy. J. Neurosurg. 19, 89– 100 (1962).

18. Agarwal, N., Choi, P. A., Shin, S. S., Hansberry, D. R. & Mammis, A. Anterior cingulotomy for intractable pain. Interdisciplinary Neurosurgery 6, 80–83 (2016).

19. Strauss, I. et al. Double anterior stereotactic cingulotomy for intractable oncological pain. Stereotact. Funct. Neurosurg. 95, 400–408 (2017).

20. Spooner, J., Yu, H., Kao, C., Sillay, K. & Konrad, P. Neuromodulation of the cingulum for neuropathic pain after spinal cord injury. Case report. J. Neurosurg. 107, 169–172 (2007).

21. Farrell, S. M., Green, A. & Aziz, T. The current state of deep brain stimulation for chronic pain and its context in other forms of neuromodulation. Brain Sci. 8, (2018).

22. Johansen, J. P., Fields, H. L. & Manning, B. H. The affective component of pain in rodents: direct evidence for a contribution of the anterior cingulate cortex. Proc Natl Acad Sci USA 98, 8077–8082 (2001).

23. Juarez-Salinas, D. L. et al. GABAergic cell transplants in the anterior cingulate cortex reduce neuropathic pain aversiveness. Brain 142, 2655–2669 (2019).

24. LaGraize, S. C. & Fuchs, P. N. GABAA but not GABAB receptors in the rostral anterior cingulate cortex selectively modulate pain-induced escape/avoidance behavior. Exp. Neurol. 204, 182–194 (2007).

25. Johansen, J. P. & Fields, H. L. Glutamatergic activation of anterior cingulate cortex produces an aversive teaching signal. Nat. Neurosci. 7, 398–403 (2004).

26. Bannister, K. et al. Multiple sites and actions of gabapentin-induced relief of ongoing experimental neuropathic pain. Pain 158, 2386–2395 (2017).

27. Donahue, R. R., LaGraize, S. C. & Fuchs, P. N. Electrolytic lesion of the anterior cingulate cortex decreases inflammatory, but not neuropathic nociceptive behavior in rats. Brain Res. 897, 131–138 (2001).

28. LaGraize, S. C., Borzan, J., Peng, Y. B. & Fuchs, P. N. Selective regulation of pain affect following activation of the opioid anterior cingulate cortex system. Exp. Neurol. 197, 22– 30 (2006).

29. Qu, C. et al. Lesion of the rostral anterior cingulate cortex eliminates the aversiveness of spontaneous neuropathic pain following partial or complete axotomy. Pain 152, 1641–1648 (2011).

30. Navratilova, E. et al. Endogenous opioid activity in the anterior cingulate cortex is required for relief of pain. J. Neurosci. 35, 7264–7271 (2015).

31. Gomtsian, L. et al. Morphine effects within the rodent anterior cingulate cortex and rostral ventromedial medulla reveal separable modulation of affective and sensory qualities of acute or chronic pain. Pain 159, 2512–2521 (2018).

32. Barthas, F. et al. The anterior cingulate cortex is a critical hub for pain-induced depression. Biol. Psychiatry 77, 236–245 (2015).

33. Fuchs, P. N., Peng, Y. B., Boyette-Davis, J. A. & Uhelski, M. L. The anterior cingulate cortex and pain processing. Front. Integr. Neurosci. 8, 35 (2014).

34. Terrasa, J. L. et al. Anterior cingulate cortex activity during rest is related to alterations in pain perception in aging. Front. Aging Neurosci. 13, 695200 (2021).

35. Kasanetz, F. & Nevian, T. Increased burst coding in deep layers of the ventral anterior cingulate cortex during neuropathic pain. Sci. Rep. 11, 24240 (2021).

36. Devinsky, O., Morrell, M. J. & Vogt, B. A. Contributions of anterior cingulate cortex to behaviour. Brain 118 (Pt 1), 279–306 (1995).

37. Deutsch, G. & Samra, S. K. Effects of nitrous oxide on global and regional cortical blood flow. Stroke 21, 1293–1298 (1990).

38. Gyulai, F. E., Firestone, L. L., Mintun, M. A. & Winter, P. M. In vivo imaging of human limbic responses to nitrous oxide inhalation. Anesth. Analg. 83, 291–298 (1996).

39. Dashdorj, N. et al. Effects of subanesthetic dose of nitrous oxide on cerebral blood flow and metabolism: a multimodal magnetic resonance imaging study in healthy volunteers. Anesthesiology 118, 577–586 (2013).

40. Reinstrup, P. et al. Regional cerebral metabolic rate (positron emission tomography) during inhalation of nitrous oxide 50% in humans. Br. J. Anaesth. 100, 66–71 (2008).

41. Ghosh, K. K. et al. Miniaturized integration of a fluorescence microscope. Nat. Methods 8, 871–878 (2011).

42. Corder, G. et al. An amygdalar neural ensemble that encodes the unpleasantness of pain. Science 363, 276–281 (2019).

43. Chen, T.-W. et al. Ultrasensitive fluorescent proteins for imaging neuronal activity. Nature 499, 295–300 (2013).

44. Zeisel, A., et al. Brain structure. Cell types in the mouse cortex and hippocampus revealed by single-cell RNA-seq. Science 347, 1138–1142 (2015).

45. Tasic, B. et al. Adult mouse cortical cell taxonomy revealed by single cell transcriptomics. Nat. Neurosci. 19, 335–346 (2016).

46. Fuzik, J. et al. Integration of electrophysiological recordings with single-cell RNA-seq data identifies neuronal subtypes. Nat. Biotechnol. 34, 175–183 (2016).

47. Harris, K. D. & Mrsic-Flogel, T. D. Cortical connectivity and sensory coding. Nature 503, 51–58 (2013).

48. Niell, C. M. & Scanziani, M. How cortical circuits implement cortical computations: mouse visual cortex as a model. Annu. Rev. Neurosci. 44, 517–546 (2021).

49. Blackwell, J. M. & Geffen, M. N. Progress and challenges for understanding the function of cortical microcircuits in auditory processing. Nat. Commun. 8, 2165 (2017).

50. Wood, K. C., Blackwell, J. M. & Geffen, M. N. Cortical inhibitory interneurons control sensory processing. Curr. Opin. Neurobiol. 46, 200–207 (2017).

51. Cichon, J., Blanck, T. J. J., Gan, W.-B. & Yang, G. Activation of cortical somatostatin interneurons prevents the development of neuropathic pain. Nat. Neurosci. 20, 1122–1132 (2017).

52. Dragunow, M. & Faull, R. The use of c-fos as a metabolic marker in neuronal pathway tracing. J. Neurosci. Methods 29, 261–265 (1989).

53. Mitchell, K. et al. Nociception and inflammatory hyperalgesia evaluated in rodents using infrared laser stimulation after Trpv1 gene knockout or resiniferatoxin lesion. Pain 155, 733–745 (2014).

54. Roome, R. B. et al. Phox2a defines a developmental origin of the anterolateral system in mice and humans. Cell Rep. 33, 108425 (2020).

55. Wheeler-Aceto, H. & Cowan, A. Standardization of the rat paw formalin test for the evaluation of analgesics. Psychopharmacology (Berl) 104, 35–44 (1991).

56. Huang, T. et al. Identifying the pathways required for coping behaviours associated with sustained pain. Nature 565, 86–90 (2019).

57. Sherrington, C. S. The integrative action of the nervous system. (Yale University Press, 1911). doi:10.1037/13798-000.

58. Bradley, N. S. & Smith, J. L. Neuromuscular patterns of stereotypic hindlimb behaviors in the first two postnatal months. III. Scratching and the paw-shake response in kittens. Brain Res. 466, 69–82 (1988).

59. Sanders, R. D., Tononi, G., Laureys, S. & Sleigh, J. W. Unresponsiveness ≠ unconsciousness. Anesthesiology 116, 946–959 (2012).

60. Sanders, R. D. et al. Incidence of Connected Consciousness after Tracheal Intubation: A Prospective, International, Multicenter Cohort Study of the Isolated Forearm Technique. Anesthesiology 126, 214–222 (2017).

61. Lennertz, R. et al. Connected consciousness after tracheal intubation in young adults: an international multicentre cohort study. Br. J. Anaesth. (2022) doi:10.1016/j.bja.2022.04.010.

62. Sebel, P. S. et al. The incidence of awareness during anesthesia: a multicenter United States study. Anesth. Analg. 99, 833–839 (2004).

63. Quiquempoix, M. et al. Layer 2/3 pyramidal neurons control the gain of cortical output. Cell Rep. 24, 2799–2807.e4 (2018).

64. Vander Weele, C. M., et al. Dopamine enhances signal-to-noise ratio in cortical-brainstem encoding of aversive stimuli. Nature 563, 397–401 (2018).

65. Fujinaga, M. & Maze, M. Neurobiology of nitrous oxide-induced antinociceptive effects. Mol. Neurobiol. 25, 167–189 (2002).

66. Homayoun, H. & Moghaddam, B. NMDA receptor hypofunction produces opposite effects on prefrontal cortex interneurons and pyramidal neurons. J. Neurosci. 27, 11496–11500 (2007).

67. Fillinger, C., Yalcin, I., Barrot, M. & Veinante, P. Afferents to anterior cingulate areas 24a and 24b and midcingulate areas 24a’ and 24b’ in the mouse. Brain Struct. Funct. 222, 1509–1532 (2017).

68. Meda, K. S. et al. Microcircuit Mechanisms through which Mediodorsal Thalamic Input to Anterior Cingulate Cortex Exacerbates Pain-Related Aversion. Neuron 102, 944–959.e3 (2019).

69. Tracey, I. & Johns, E. The pain matrix: reloaded or reborn as we image tonic pain using arterial spin labelling. Pain 148, 359–360 (2010).

70. Peeters, L. M. et al. Chemogenetic silencing of neurons in the mouse anterior cingulate area modulates neuronal activity and functional connectivity. Neuroimage 220, 117088 (2020).

71. Basbaum, A. I. & Fields, H. L. Endogenous pain control systems: brainstem spinal pathways and endorphin circuitry. Annu. Rev. Neurosci. 7, 309–338 (1984).

72. Fillinger, C., Yalcin, I., Barrot, M. & Veinante, P. Efferents of anterior cingulate areas 24a and 24b and midcingulate areas 24a’ and 24b’ in the mouse. Brain Struct. Funct. 223, 1747–1778 (2018).

73. Linassi, F., Zanatta, P., Tellaroli, P., Ori, C. & Carron, M. Isolated forearm technique: a meta-analysis of connected consciousness during different general anaesthesia regimens. Br. J. Anaesth. 121, 198–209 (2018).

74. Gaskell, A. L. et al. Frontal alpha-delta EEG does not preclude volitional response during anaesthesia: prospective cohort study of the isolated forearm technique. Br. J. Anaesth. 119, 664–673 (2017).

75. Ní Mhuircheartaigh, R., Warnaby, C., Rogers, R., Jbabdi, S. & Tracey, I. Slow-wave activity saturation and thalamocortical isolation during propofol anesthesia in humans. Sci. Transl. Med. 5, 208ra148 (2013).

76. Pawar, N. & Barreto Chang, O. L. Burst suppression during general anesthesia and postoperative outcomes: mini review. Front. Syst. Neurosci. 15, 767489 (2021).

77. García, P. S., Kreuzer, M., Hight, D. & Sleigh, J. W. Effects of noxious stimulation on the electroencephalogram during general anaesthesia: a narrative review and approach to analgesic titration. Br. J. Anaesth. 126, 445–457 (2021).

78. Vong, L. et al. Leptin action on GABAergic neurons prevents obesity and reduces inhibitory tone to POMC neurons. Neuron 71, 142–154 (2011).

79. Hippenmeyer, S. et al. A developmental switch in the response of DRG neurons to ETS transcription factor signaling. PLoS Biol. 3, e159 (2005).

80. Taniguchi, H. et al. A resource of Cre driver lines for genetic targeting of GABAergic neurons in cerebral cortex. Neuron 71, 995–1013 (2011).

81. Tamamaki, N. et al. Green fluorescent protein expression and colocalization with calretinin, parvalbumin, and somatostatin in the GAD67-GFP knock-in mouse. J. Comp. Neurol. 467, 60–79 (2003).

82. Jackson, J., Karnani, M. M., Zemelman, B. V., Burdakov, D. & Lee, A. K. Inhibitory control of prefrontal cortex by the claustrum. Neuron 99, 1029–1039.e4 (2018).

83. Alkire, M. T. & Gorski, L. A. Relative amnesic potency of five inhalational anesthetics follows the Meyer-Overton rule. Anesthesiology 101, 417–429 (2004).

84. Dwyer, R., Bennett, H. L., Eger, E. I. & Heilbron, D. Effects of isoflurane and nitrous oxide in subanesthetic concentrations on memory and responsiveness in volunteers. Anesthesiology 77, 888–898 (1992).

85. Eger, E. I. Age, minimum alveolar anesthetic concentration, and minimum alveolar anesthetic concentration-awake. Anesth. Analg. 93, 947–953 (2001).

86. Karippacheril, J. G. & Ho, T. Y. Data acquisition from S/5 GE Datex anesthesia monitor using VSCapture: An open source.NET/Mono tool. J. Anaesthesiol. Clin. Pharmacol. 29, 423–424 (2013).

87. Zhang, J., Cavanaugh, D. J., Nemenov, M. I. & Basbaum, A. I. The modality-specific contribution of peptidergic and non-peptidergic nociceptors is manifest at the level of dorsal horn nociresponsive neurons. J Physiol (Lond) 591, 1097–1110 (2013).

88. Wasilczuk, A. Z., Meng, Q. C. & McKinstry-Wu, A. R. Electroencephalographic evidence for individual neural inertia in mice that decreases with time. Front. Syst. Neurosci. 15, 787612 (2021).

89. Jarvie, B. C., Chen, J. Y., King, H. O. & Palmiter, R. D. Satb2 neurons in the parabrachial nucleus mediate taste perception. Nat. Commun. 12, 224 (2021).

90. Mukamel, E. A., Nimmerjahn, A. & Schnitzer, M. J. Automated analysis of cellular signals from large-scale calcium imaging data. Neuron 63, 747–760 (2009).

