## Supplemental Figures for "Paradoxical increases in anterior cingulate cortex activity during nitrous oxide-induced analgesia reveal a signature of pain affect"

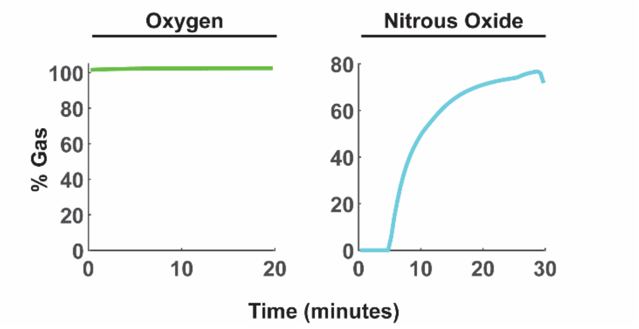


**Supplemental Fig. 1. Oxygen and nitrous oxide concentrations during recordings of spontaneous ACC activity (See Figures 1 and 2).** Gas concentrations averaged across all mice in figures 1 and 2 (mean ± SEM; N = 30 mice).

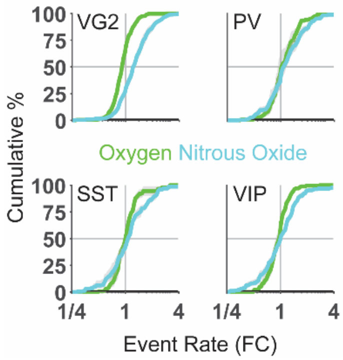


**Supplemental Fig. 2. Nitrous oxide differentially influences molecularly distinct subpopulations of ACC neurons.** Fold change in event rate quantified as cumulative percent (mean ± SEM) for distinct populations of ACC neurons (neurons expressing vGluT2 (VG2), parvalbumin (PV), somatostatin (SST), or vasoactive intestinal peptide (VIP). N = 6 mice per genotype.

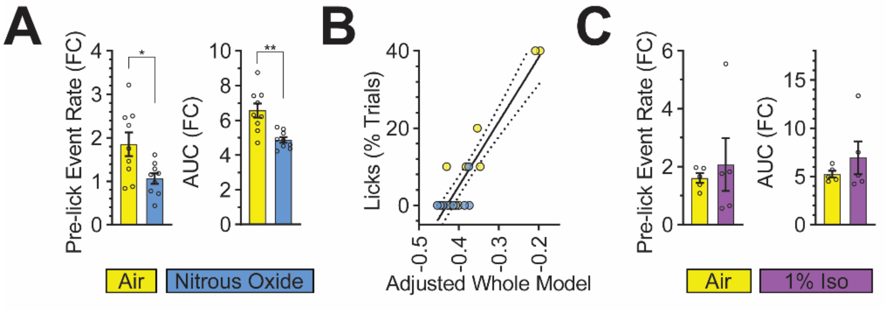


**Supplemental Fig. 3. ACC neural activity correlates with production of noxious stimulus-evoked affective-motivational behaviors.** (**A**) Nitrous oxide-induced changes to event rate post laser and prior to licking; area under the curve (AUC) quantified from **Fig. 4H** (paired t-test, n = 9). (**B**) Stepwise linear regression with interaction effects of the Adjusted Whole Model (pre-lick event rate (FC), maximum event rate (FC), AUC (FC), and percentage of laser responsive neurons vs percentage of licks; adjusted $R^2 = 0.736$, $p < 0.004$). (**C**) Isoflurane-induced changes to event rate prior to licking and AUC quantified from **Fig. 5F** (paired t-test, n = 5).
